# miR-210 locus deletion disrupts cellular homeostasis; an integrated genetic study

**DOI:** 10.1101/2024.10.18.619004

**Authors:** Mihai Bogdan Preda, Evelyn Gabriela Rusu-Nastase, Carmen Alexandra Neculachi, Xiaoling Zhong, Christine Voellenkle, Nathalie M. Mazure, Ovidiu Balacescu, Cristina Ivan, Xiao-Wei Zheng, Mihaela Gherghiceanu, Kevin Lebrigand, Maya Simionescu, Fabio Martelli, Bernard Mari, Sergiu-Bogdan Catrina, Alexandrina Burlacu, Mircea Ivan

**Author notes:** Department of Surgery, Indiana University School of Medicine, Indianapolis, IN 46202, USA. Caris Life Sciences, Irving, TX, 75039, USA.

## Abstract

MiR-210 is widely recognized as the quintessential hypoxia-responsive miRNA and thought to fine-tune various facets of cellular homeostasis. We hereby present an integrative appraisal of phenotypic and molecular repercussions of disrupting the corresponding locus in human and mouse cells using multiple genetic strategies. Briefly, MIR210 deletion led to decreased cellular fitness and suboptimal responses to several stress types. Transcriptomic comparisons using different profiling platforms, performed independently by members of this collaboration, revealed consistent deregulation of neighboring genes, in locus-disrupted cells. Interestingly, the anticipated enrichment in miR-210 targets failed to materialize in unbiased analyses. Our results point to the biological significance of unrecognized regulatory elements that overlap miRNA genes and should serve as note of caution for studies based for genetic disruption of such loci.

## Introduction

MicroRNAs (miRNAs) are arguably the most extensively studied non-coding RNA family, thought to fine tune virtually all signaling pathways and homeostatic processes (1, 2). Dysregulation of miRNA expression is extensively documented in every major disease, including cancer, cardiovascular and neurological disorders (3). The mechanistic paradigm of miRNA action states that they regulate gene expression predominantly at posttranslational level by targeting messenger RNAs (mRNAs) for degradation, translational repression or a combination thereof (4–6).

Despite over two decades of studies and more than a million indexed publications focusing on, or containing miRNA, major dilemmas persist in the field. For example, while each miRNA is suspected to regulate multiple genes, the exact number of biologically relevant targets remains unclear. To further compound the controversy, the term “confirmed targets” is used too liberally, without solid evidence that the corresponding interactions actually occurs in vivo (7).

While most microRNA experimental studies have relied on reagents that either mimic or inhibit the major single-stranded RNA product (8–10), genetic interventions on these loci are thought to provide powerful biological insights (11). A major potential flaw in interpreting such experiments is that the outcomes are typically assessed solely through the lens of targets, with rare or no consideration given to the genomic context. The case of miR-210 exemplifies this challenge. Whereas we have a relatively good understanding of its regulatory inputs, such as oxygen deprivation, immune and inflammatory signals (9, 10, 12, 13), the mechanisms responsible for its actions remain controversial. An in-depth examination of the available literature reveals a heavy bias for small sets or even unique targets, including ISCU, NDUFA4, SDHD, HIF1A (8, 14, 15). However, an exhaustive list of miR-210 candidate targets using the available bioinformatic resources can make the case for hundreds, if not thousands, potential genes (16). Furthermore, it is unclear to what extent genes regulated by mature miR-210 remain relevant in knockout models, as published studies often overlook this issue or approach it with a bias towards a limited number of candidates. Of note, these dilemmas are highly relevant for the miRNA field as a whole.

We present a multicenter study integrating genetic interventions in mouse and human cells with pertinent insights from large OMICS datasets. Our investigation provides new evidence of MIR210 locus roles in cellular homeostasis and stress responses and emphasizes the importance of considering overlapping regulatory elements for a rigorous understanding of miRNA loci.

## Result

### Genetic inactivation of MIR210 locus in human and mouse cells

The human MIR210 gene, first identified as hypoxia-inducible through miRNA microarray analysis (17), is situated within an intron of a lncRNA gene known as the MIR210 host gene (MIR210HG) in chromosome 11p15.5 region. MIR210 knockout (KO) HEK cells were generated by deleting ∼30 nucleotides with a bidirectional CRISPR-Cas9 approach. Using a similar strategy, we also engineered an ∼2.5kb deletion that eliminates the entire MIR210HG sequence, including the intronic MIR210 locus (Fig. 1a, Suppl Fig. 1). Homozygous knockout clones (MIR210^-/-^ and MIR210HG^-/-^) were identified by genomic PCR (Fig. 1b) and expanded for subsequent studies. Loss of RNA products was confirmed by RT-qPCR analysis of miR-210 and MIR210HG (exon 2 transcript) under both high and low oxygenation (Fig. 1c-d).

**Figure 1.**
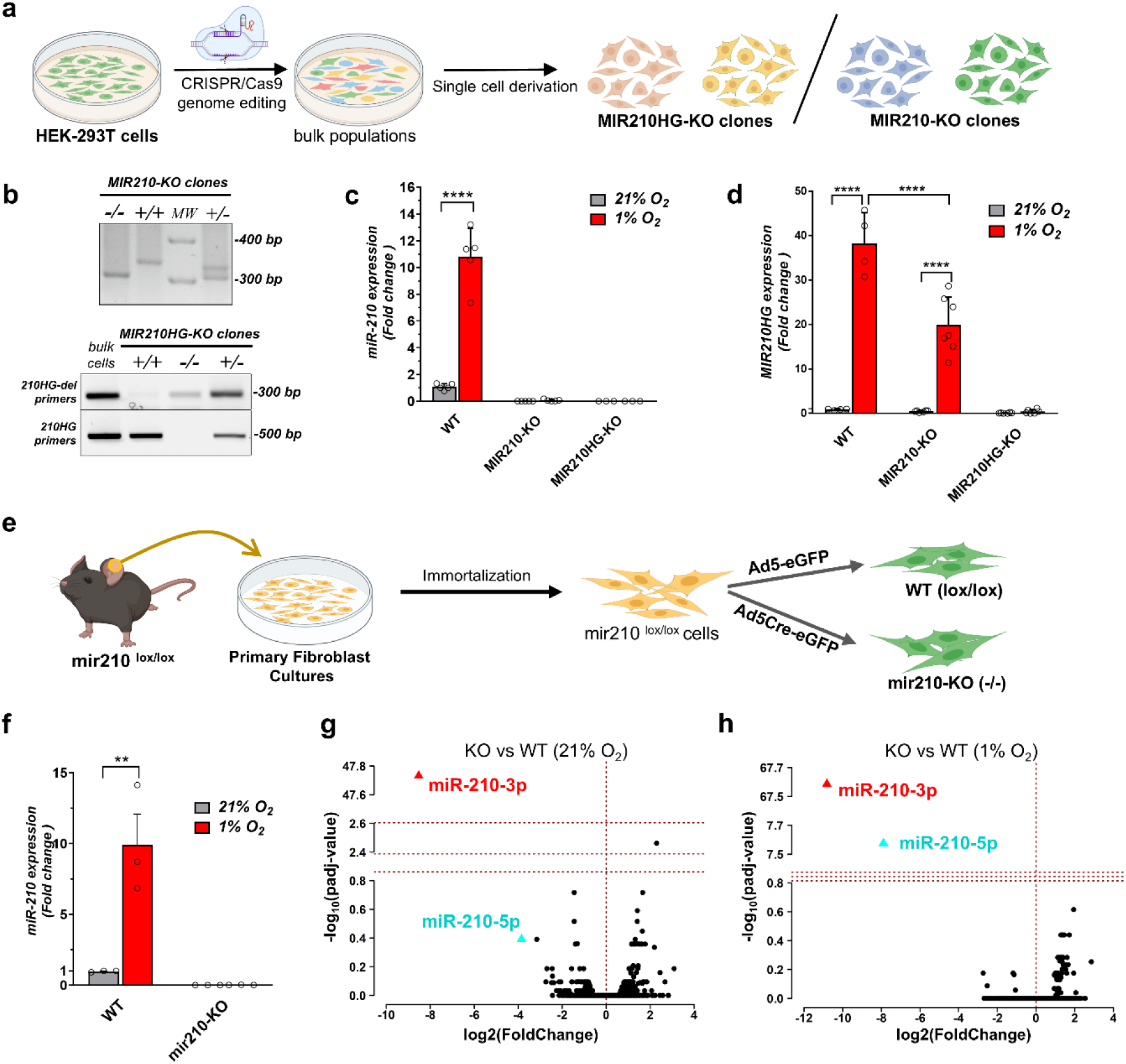
Generation and validation of experimental models. (a) Schematic overview illustrating the generation of HEK knockout (KO) cell clones via CRISPR-Cas9 genome editing, created using BioRender.com (https://biorender.com/); (b) Genomic validation by PCR analysis confirms single-cell-derived KO clones for MIR210 or MIR210HG, identifying both heterozygous (+/-) and homozygous (-/-) genomic deletions. (c-d) Relative levels of miR-210-3p, and MIR210HG transcripts in wild-type (WT), MIR210–KO, and MIR210HG–KO cells under 21% or 1% O_2_ for 24 hours. Data shown as mean ± SD and the statistical significance determined by two-way ANOVA (****p < 0.0001). (e) Experimental workflow of mir210 locus excision in immortalized primary fibroblasts from mir210 lox/lox male mice, created with BioRender.com (https://biorender.com/); (f) Data are shown as mean ± SD, with statistical significance determined by two-way ANOVA (**p < 0.01) (g-h) Volcano plots showing differential expression of miRNAs in mir210-KO mouse fibroblasts under 21% or 1% O_2_, with significantly downregulation of miR-210-5p and miR-210-3p.

Mouse immortalized fibroblasts were derived from male mir210^lox/lox^ mice (11, 13, 18). Briefly, dermal fibroblasts were isolated from adult mouse ears and immortalized by shRNA-mediated p53 inactivation. Acute excision of mir210 was achieved by infection with Ad5Cre-eGFP versus Ad5eGFP-only control (Fig. 1e) and confirmed by RT-qPCR for miR210-3p (Fig. 1f). Reassuringly, in small RNA-seq profiling miR-210-3p and -5p were among the top mature miRNAs differentially expressed between WT *versus* KO cells (Fig.1g-h).

### Phenotypic repercussions of MIR210 locus loss

The relative fitness of locus-deleted and wild type cells was investigated in competition assays (Fig. 2a). The results showed that WT cells consistently outperformed KOs (Fig. 2a), with similar advantage over both small and large deletion. This is consistent with the presence of MIR210HG among the 3% of lncRNA loci important for cell growth (19). In cellular respirometry and metabolic stress assays using the Seahorse XFp system MIR210-deficient populations exhibit an abnormal dynamic of OCR shift (Fig. 2b, d). Deletion also led to mild, but consistent alteration of cell cycle dynamic following oxygen deprivation (Fig. 2c, f). In the context of HEK cells MIR210-KOs are less responsive to hypoxia than wild type controls from the standpoint of S and G2/M phase accumulation (Fig. 2c). A similar effect was observed following the more extensive deletion involving the entire MIR210HG (Suppl Fig. 2). Furthermore, at least in murine fibroblasts, locus deletion is followed by excessive mitochondrial ROS production (Fig. 2e) and altered mitochondrial volume particularly under oxygen deprivation stress (Fig. 2g).

**Figure 2.**
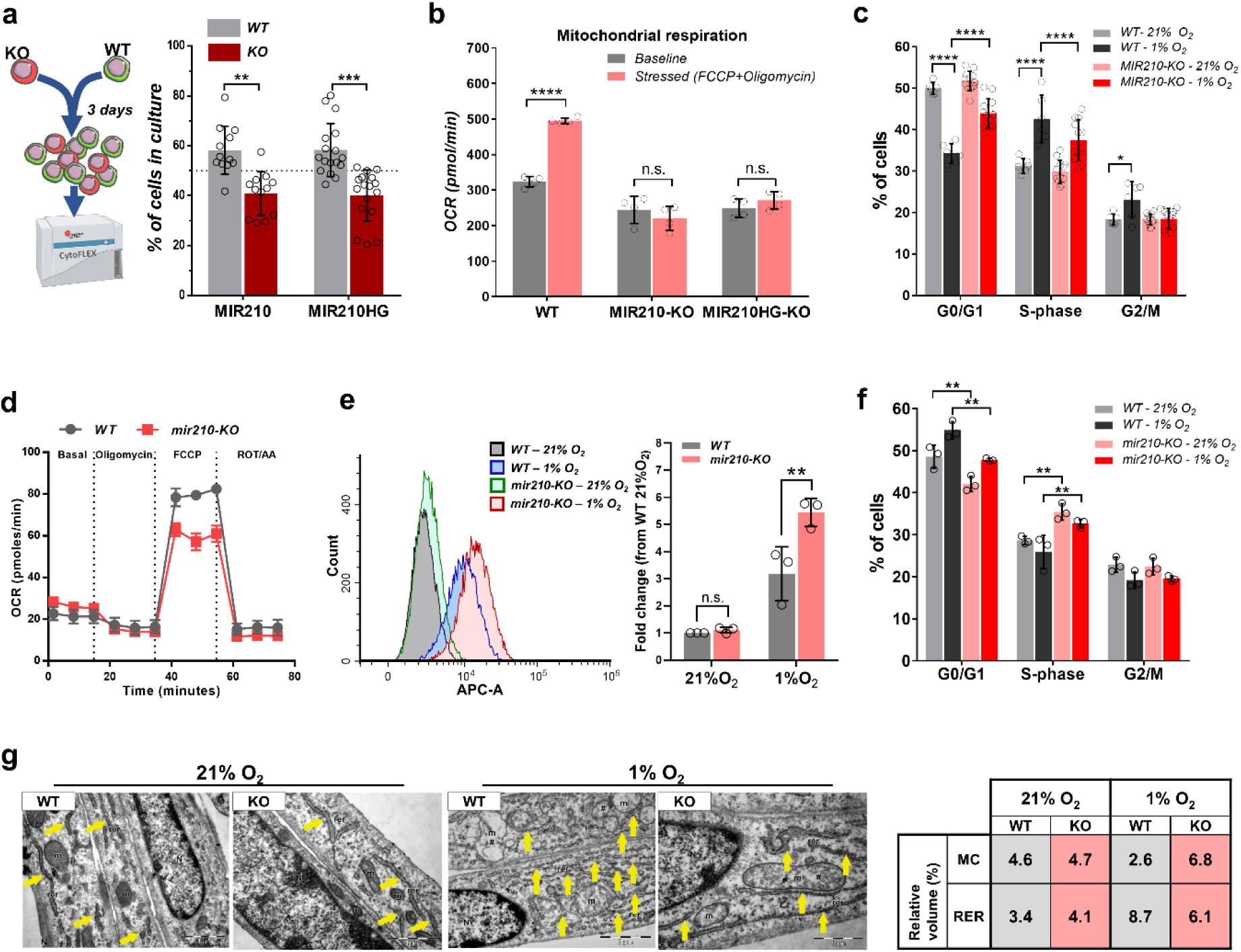
Phenotypic and metabolic impact of MIR210 locus deletion. (a) Competitive growth assays of WT and MIR210 or MIR210HG KO clones labeled with GFP or dTomato, analyzed after 3 days of coculture. Flow cytometry results show significantly fewer KO cells compared to WT (mean ± SD from 3 experiments, with four different clones analyzed per experiment; two-way ANOVA, **p < 0.01, ***p < 0.001). (b) Mitochondrial respiration (OCR) in HEK WT and KO clones under basal or stress conditions. Data are mean ± SD from 2 experiments in duplicate (two-way ANOVA, n.s. not significant, ***p < 0.001, ****p < 0.0001). (c) Significant changes in cell cycle distribution in HEK cells (WT and MIR210-KO) under different oxygen levels (two-way ANOVA, *p < 0.05, ****p < 0.0001). (d) OCR profiles from a Mito Stress Test in WT and mir210-KO mouse fibroblasts reveal reduced maximal respiration and spare capacity in KO cells (mean ± SD from a single representative experiment, three replicates). (e) Mitochondrial ROS quantification in WT and mir210-KO mouse fibroblasts after 24-hour exposure using MitoSOX and flow cytometry (mean ± SD from 3 experiments, two-way ANOVA, **p < 0.01). (f) Cell cycle alterations in mir210-KO mouse fibroblasts under different oxygen levels (two-way ANOVA, **p < 0.01). (g) Cellular ultrastructure by transmission electron microscopy shows structural variations in mitochondria and endoplasmic reticulum in KO fibroblasts under hypoxic conditions.

Collectively, these experiments indicate that cellular homeostasis and at least some adaptive responses to stress are subtly but consistently altered as a result of eliminating MIR210 locus. Persistent cellular dysfunction of this magnitude may be particularly relevant for specific chronic diseases, that often take decades to manifest.

### Consequences of miR-210 disruption on molecular programs

We performed transcriptomic profiling of mouse and human cells in low and high oxygen conditions and pathway enrichment analysis was performed using publicly available tools: Enrichr (https://maayanlab.cloud/Enrichr/) and MSigDB hallmark gene sets to capture specific biological states. Results are summarized in Fig. 3 and Supplemental Fig. 3. Among the top hallmark differences between KO and WT are hypoxia and mTORC1 signaling (Fig. 3a-b) as well as cycle related signatures (E2F targets, G2-M checkpoint) (Fig. 3c-f). Additionally, glycolysis and mitochondrial OXPHOS were identified among the top differential metabolic signatures.

**Figure 3.**
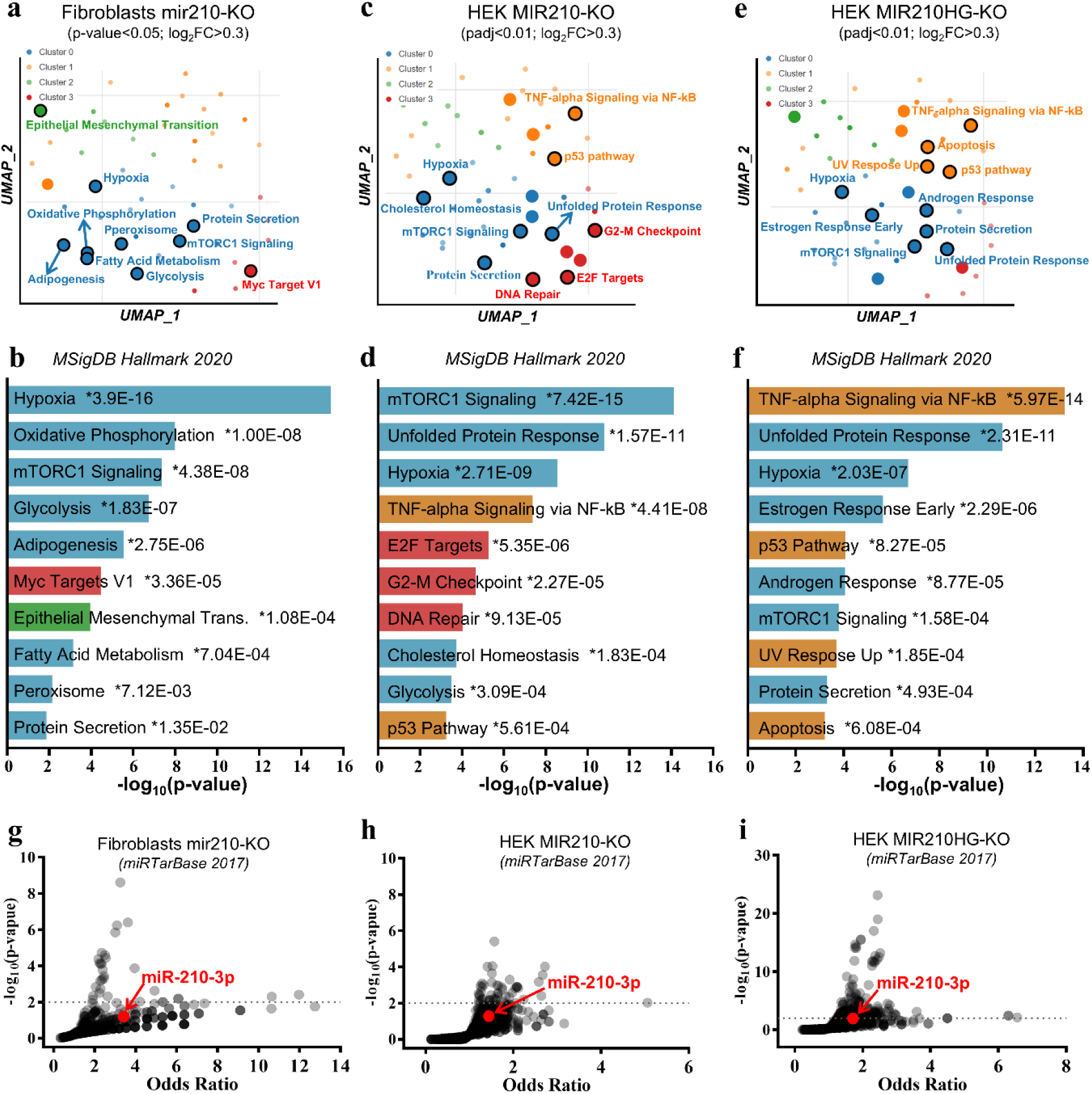
Transcriptomic consequences of locus disruption in mouse and human cells. Pathway enrichment analysis of gene expression in miR210-KO mouse fibroblasts and HEK cells under low O_2_ (1%) was performed using Enrichr and visualized with an Appyter for bulk RNA-seq (https://appyters.maayanlab.cloud/#/Bulk_RNA_seq). (a) Scatter plot with larger, black-outlined points highlighting significantly enriched pathways, grouped by gene set similarity. (b) Bar charts displaying the most significant gene sets from MSigDB_Hallmark_2020, with levels of significance indicated. (c) Volcano plots showing microRNAs potentially affecting downregulated DEGs in mir210-KO cells, with each microRNA represented by its odds ratio and statistical significance. (d-f) Similar analysis for MIR210-KO HEK cells under 1% O_2_, applying a stricter significance threshold (p-adjusted < 0.01) to limit gene number for more precise enrichment analysis. (g-i) Parallel analysis in MIR210HG-KO HEK cells under similar experimental setups, detailing transcriptomic shifts and pathway dynamics.

While bioinformatic analyses list tentative directional changes for specific signatures, we caution against a simplistic interpretation that the respective pathways are overactive or conversely inhibited after miR210 locus loss. In our view, a more prudent interpretation is that KO cell responses to stress, including hypoxia, are suboptimally coordinated. For example, while the canonical hypoxic response is still present in KO cells, some HIF targets are excessively induced (SLC2A1, VEGFA, PGK1, NOS2) whereas others respond less or not at all compared to WT cells (CA9, GAPDH, EGLN3), at least at the time of investigation. Overall, our analysis is consistent with a suboptimal response to changes in ambient oxygen following MIR210 locus disruption.

We next directed our attention to miR-210 targets, which remains a controversial topic in studies based on miRNA deletion models. It is noteworthy that the number of miR-210 target candidates amounts to a significant fraction of all coding genes when multiple prediction programs are queried. As such, claims of mechanistic targets can be readily made based on convenient choices from differentially expressed genes (DEG) in KO versus WT. Published studies rarely present the outcome of miRNA KOs tested in unbiased target enrichment analyses.

Specifically for our case, if mature miR-210 were the most important element eliminated by deletion, its targets would stand out in enrichment analyses particularly in hypoxia. Contrary to this expectation, in Enrichr analysis using the miRTarBase computational prediction of miRNA-target gene interaction, miR-210 targets are collectively „lost” in a cloud of miRNAs, without significant enrichment in mouse or human KOs (Fig 3g-i). Furthermore, widely validated targets such as ISCU, EFNA3, HOXA3 fail to exhibit the expected pattern: higher expression in KO, particularly in hypoxia. Arguably the most relevant is ISCU, with little doubt a bona fide target of miR-210, as its interaction with mir210 is listed in high throughput RNA-RNA interaction datasets in both human and mouse cells (http://bigdata.ibp.ac.cn/npinter5/). Taken together, these results suggest that the biochemical and phenotypical features of KO cells cannot be automatically attributed to miR-210 regulation of targets.

These observations suggests that our interventions, or indeed similar strategies leading to miRNA KO models, are prone to affect a multitude of genomic features and as such exhibit phenotypes not completely attributable to the elimination of mature miRNA product. Unbiased information on genetic elements and chromatin architecture contained in large public datasets provides important insights into the transcriptional landscape of genetically modified cells. To this end, we examined in detail the genomic neighborhood around MIR210, a region characterized by high synteny and evolutionary conservation (Suppl Fig. 4a-b).

Indeed, MIR210/MIR210HG locus exhibits extensive long-range chromatin interactions with nearby loci in a variety of human cells, including HEK293 (Fig. 4c; Hi-C data from 3DIV database, http://3div.kr).

**Figure 4.**
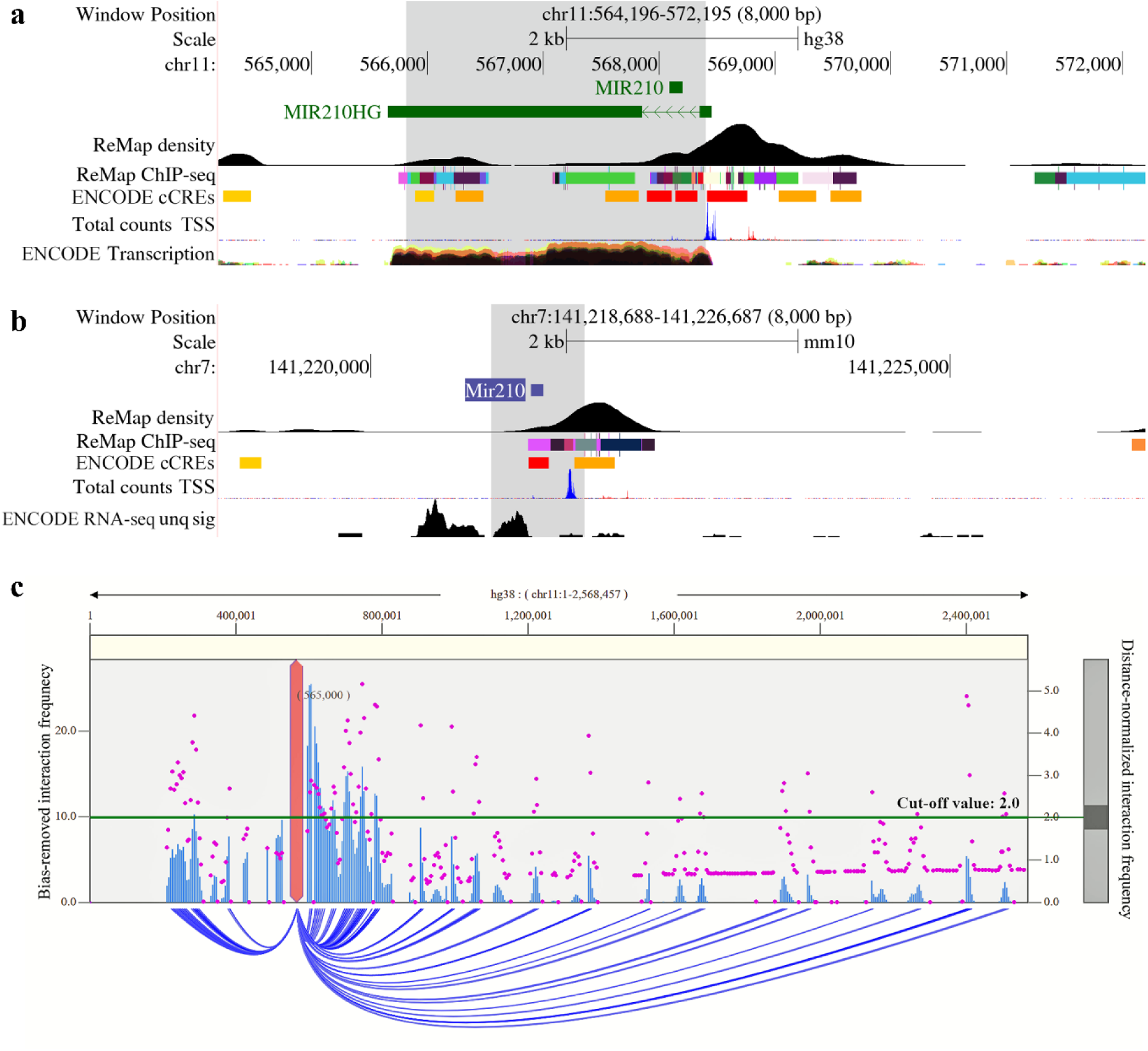
Genomic Features of human (a,c) and mouse (b) miR210 locus. ENCODE database summary of transcript abundance, transcriptional regulator binding (ReMap density, ChIP-seq), chromatin states (red: active promoters; orange: enhancers) and mapped transcription start sites (TSS, FANTOM5). (c) Hi-C interactions in HEK293T cells anchored on MIR210HG locus (3DIVdatabase). Interaction frequency is represented by magenta dots and blue arcs correspond to interactions above the cut-off value (green line).

In locus-deleted cells neighborhood genes appear broadly deregulated in transcriptomics data with changes varying depending on deletion size and oxygen availability (Fig. 5a-b, Suppl Fig. 5a). Further, time-course RT-qPCR assays in synchronized populations confirmed the broad downregulaton of neighborhood genes (*RNH1*, *HRAS*, *RASSF7*, *PHRF1*, *IRF7*, *DEAF1*, *TMEM80*, *GATD1*, and *CD151*) particularly in hypoxia, with the possible exception of *LRCC56* and *LMNTD2* (Fig. 5c). In mouse fibroblasts, the dynamic nature of the neighborhood perturbations was notable (Suppl Fig. 5b), supporting the need to consider the time component when investigating the effect of eliminating loci suspected to play roles in feedback responses. It is plausible that deregulation of a multitude of coding, non-target, genes could alone explain the masking of a conventional miRNA effects.

**Figure 5.**
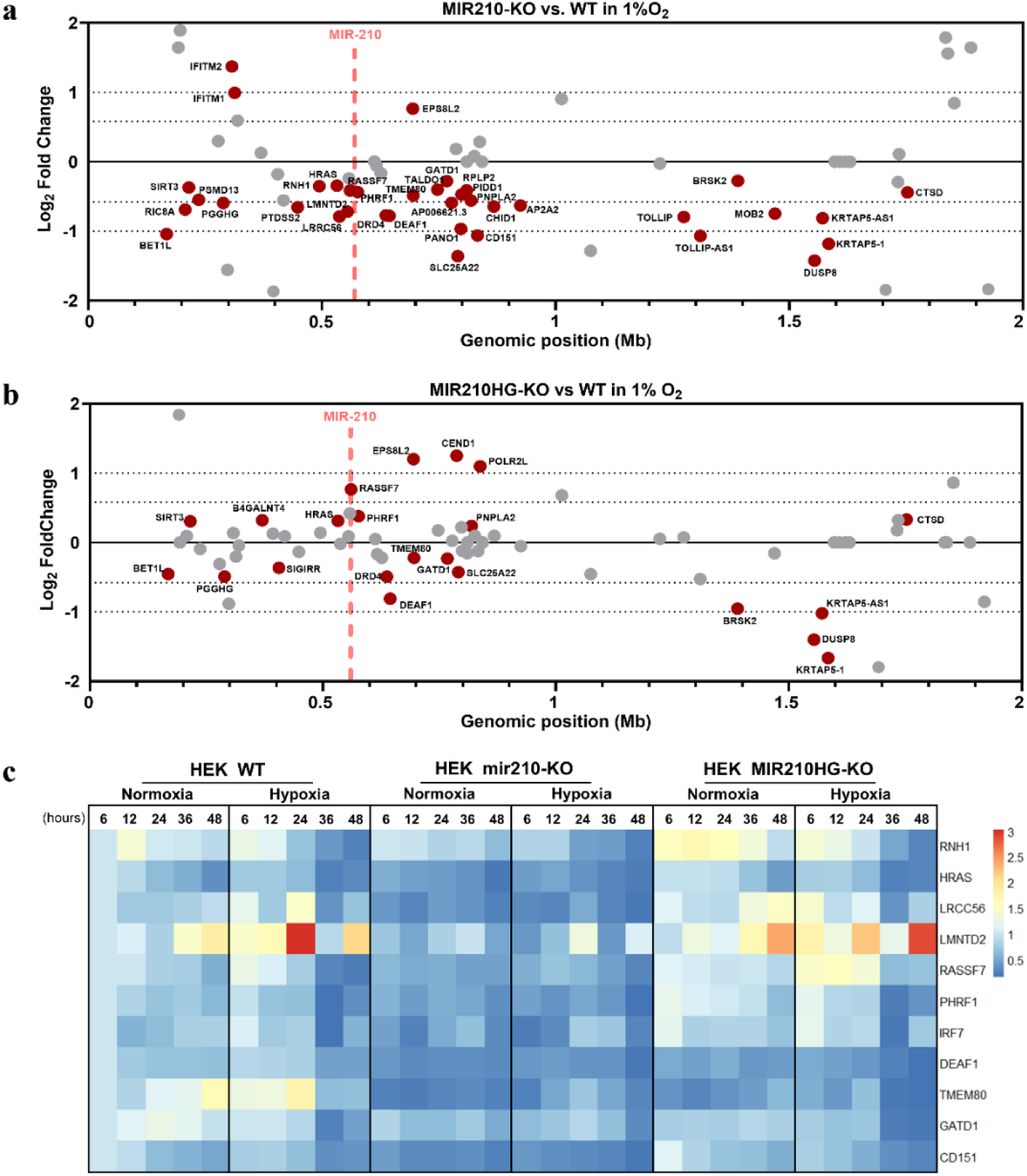
Effect of MIR210 locus deletion on local transcriptional gene regulation in human cells. Summary of local transcriptional changes upon MIR210 (a) or MIR210HG (b) deletion in human cells (HEK293T) exposed to low O_2_ (1%) for 24 hours, versus wild-type cells. Genes are plotted based on their chromosomal position on the x axis and their differential expression (log2FoldChange) on the y-axis. Genes significantly differentially expressed (padj < 0.05) were name-labelled and marked with red dots. MIR210 genomic locus is highlighted by a vertical dashed line. (c) Heatmap visualization of qRT-PCR analysis of specified neighboring genes at various time points in both 21% and 1% O_2_. Data are presented as fold changes relative to HEK WT Normoxia at 6 hours and are normalized on a per-row color scale for clarity.

We should also consider another potentially significant element affected by our genetic intervention, MIR210HG, a lncRNA relatively abundant in some cell types. This poorly characterized transcript is completely absent in the cells harboring the large deletion and also significantly downregulated in the clones with the 30-nt cut. Furthermore, although neither mm10 or mm39 genomes list a murine MIR210HG ortholog, its existence is plausible based on previous research (20), and information from public datasets: cis-regulatory elements (cCREs), RNAseq signal and sequence homology (Fig. 4a-b). Due to the very nature of how the mouse KO was generated, the putative murine lncRNA would also have been eliminated following AdenoCre mediated excision. Overall, inadvertent or unavoidable lncRNA elimination (partial or total) needs to be considered as a potential contributor to the biochemical or phenotypical outcome.

Messages consistent with our results are conveyed by naturally occurring variations in human populations and their correlations with gene expression extracted from Genotype-Tissue Expression (GTEx) database (21). As summarized in Suppl Fig. 6a-b local single nucleotide polymorphisms are almost exclusively associated with variations in the expression of neighborhood genes (rather than MIR210HG itself), at least in some tissues. From this perspective, it only makes sense that comparatively dramatic genomic alterations, such as our engineered deletions, should also be consequential for a multitude of neighborhood genes. Such effects, however, should not be considered as an automatic feature of interference (whether natural or experimental) with miRNA loci. As illustrated in Suppl Fig 7, naturally occurring variants at another hypoxia-responsive intergenic noncoding locus, miR193BHG/lincNORS (22), appear particularly significant for the lncRNA itself, but less so for the abundance of neighboring transcripts.

Taken together, the above results indicate that the impact of MIR210 locus cannot be simplistically reduced to a handful of (conveniently chosen) targets of mature miR-210-3p. Indeed, its relevance for cellular homeostasis and stress responses needs to be understood in a cell-specific context as the combined activity of RNA products (miRNA & lncRNA) and regulatory DNA elements.

## Discussion

To address long standing dilemmas in miRNA biology, we conducted a comprehensive investigation using genetically engineered human and murine cells. Despite a vast, and still expanding, body of literature on this subject, the depth of our understanding of miRNA biology has been increasingly questioned. Persistent misconceptions and misunderstandings have impacted the overall quality of many publications on this topic that appear in PubMed each year (23). Setting aside the unknown negative impact of papermills (24, 25), most miRNA studies rely on exogenous transduction of reagents (such as “mimic” or “anti-miR”). The specificity of these reagents is often assumed without rigorous testing and there is frequently minor consideration of the natural abundance of miRNAs. Comprehensive analysis of miR-210 target candidates predicted by various large-scale biological databases for miRNA-target interactions often identify a significant proportion of genes in mammalian genomes, which is biologically improbable. Furthermore, claims of previously validated targets should be interpreted with caution.

Additional complexities arise when comparing the above experimental strategies with targeting miRNAs at genomic level using Cre-Lox or CRISPR – based interventions (26–28). Typically, these studies report subtle phenotypes, in contrast to the more pronounced effects observed with mimic and anti-miRNAs, raising concerns that the latter systems may be tested outside of physiologic range. Mechanistically, the relative contributions of individual targets to the observed phenotypes or their impact on gene expression in KO models are often overestimated. There is a notable lack of publications demonstrating that the targets of a given miRNA stand out collectively in unbiased analyses.

Our study demonstrates that the loss of miR-210 locus leads to reduced cellular fitness, a finding consistent with previous experiments involving miR-210 antagonists (8–10). However, it seems unlikely that the phenotype results solely from the absence of mature miR-210, as our data did not reveal a specific target signature. More critically, we noted deregulation in the expression of neighboring coding genes, some of which are highly expressed. Unfortunately, our findings suggests that it is not feasible to fully rescue the expression and phenotype, as restoring the dynamic expression of these neighboring genes to reproduce the WT response may be impossible.

Relevant for our study, it is important to note that miRNA KO studies often pay limited attention to the regulatory elements disrupted by the intervention, as well as the effects on neighboring genes and potential phenotypic or biochemical ramifications (13, 18, 29–31). In contrast, our study highlights that altered expression of neighboring gene represents a major theme, and the specific miR-210 target signature was weak or absent. It is noteworthy that MIR210 genomic locus engages in numerous chromatin interactions with neighboring genes, including RASSF7, HRAS1, PHRF1, TMEM80, RNH1. It stands to reason that disruption of the locus may impact some of these genes, but the experiments provide valuable insights into the specific genes involved, the magnitude of the impact, and the dynamic nature of these effects. Databases such as GTEx and Dice provide potential clues that neighboring genes may be particularly sensitive to genetic changes in the MIR210HG/miR-210 locus. Therefore, it is reasonable to assume that a more substantial intervention at this location (i.e., Cre-Lox or CRISPR cuts) could significantly deregulate neighboring genes, potentially blurring or even erasing the anticipated miRNA-specific signature.

Last but not least, many miRNAs are located within introns of lncRNA genes, and targeting the miRNAs may inadvertently affect the expression of the lncRNAs. Notably, in the GTEx database, MIR210HG shows similar or higher expression compared to lncRNAs with established biological activity (Suppl Fig. 8) and its disruption may therefore have significant biochemical or phenotypical consequences.

appears similar or highly expressed compared to lncRNAs with established biological activity Integrated experimental models, like those described above, are likely to become increasingly relevant in the future. Numerous copy number variations, including relatively small deletions affecting miR-210 locus, have been cataloged in human populations and are often listed as having “unknown significance”. The region containing miR-210 on Chr11p15.5 is generally overshadowed by the more effectively studied centromeric segment, which includes IGF2/H19 imprinted locus (32, 33). Currently, single nucleotide variations at MIR210 genomic locus, including adjacent genes, are linked to chronic conditions, notably Systemic Lupus erythematosus (SLE) (34, 35) aligning with the immune and inflammatory abnormalities observed in mir210-KO mice (13, 30). Our study underscores the need for further exploration of this genomic region, which harbors a wealth of underexplored coding and regulatory elements pertinent to both physiology and disease. While the data should, at a minimum, prompt the need to revisit miR-210 knockout studies, our investigation is likely more broadly informative for the biology of miRNA loci.

## Methods

### Generation of mir210 KO mouse fibroblasts

C57BL/6 mir210 floxed mice were obtained by M. Ivan (Indiana University School of Medicine, Indianapolis, IN) in collaboration with N. Bardeesy (Massachusetts General Hospital, Boston, MA), as previously described (18). Housing and all experimental animal procedures followed the NIH research guidelines and were approved by the Institutional Animal Care and Use Committee of the Indiana University School of Medicine.

Primary dermal fibroblasts were isolated from the ears of mir210 floxed male mice (8–12-week-old), as described (18). Two passages later cells subjected to inactivation of p53 using an shRNA-expressing lentivirus followed by puromycin (2ug/ml) selection, a well-established immortalization method. Locus deletion was next achieved by Ad5CMVCre-eGFP Adenovirus infection, as described (13). Ad5CMV-eGFP infection was performed on control cells. Deletion of miR210 was confirmed by genomic PCR and absence of mature mmu-miR-210-3p by TaqMan MicroRNA assay (A25576, ThermoFisher Scientific) and TaqMan MicroRNA Reverse Transcription Kit (4366596, ThermoFisher Scientific). U6 expression was quantified for data normalization.

### CRISPR-Cas9 mediated deletions

HEK293T cell line purchased from the American Type Culture Collection (ATCC) was used to generate human KO cells for either MIR210 or MIR210HG. Plasmid construction and the design of the sgRNAs used to delete the selected fragments were conducted in collaboration with Transposagen Biopharmaceuticals (Lexington, United States). WT HEK cells were seeded in 6-well plates at a density of 50,000 cells per cm². The following day, cells were transfected using Lipofectamine 3000 (L3000015, ThermoFisher Scientific) in OptiMEM reduced serum medium (11058021, ThermoFisher Scientific), with two vectors containing nucleases targeting the 5’ and 3’ ends of the projected fragment for each KO type (MIR210 or MIR210HG). Specific sequences of the gRNAs are as follows: MIR210 5’ region (5’-GCGCAGTGTGCGGTGGGCAG-3’), MIR210 3’ region (5’-GCCGCTGTCACACGCACAGT-3’), MIR210HG 5’ region (5’-GCTGGCACCCTCTCGCCCCC-3’) and MIR210HG 3’ region (5’-GGGCAAGGAAGCCATCCACA -3’). Fifteen hours later, the transfection medium was replaced with growth medium, and upon reaching confluence, the cells were passaged and sampled for genomic DNA isolation to confirm deletions. Two rounds of transfection were performed, followed by the derivation of single-cell clones, which were then expanded and screened for deletion status using specific genomic PCR primers to distinguish between heterozygous and homozygous deletions.

### Cell culture and reagents

Mouse fibroblasts were cultured in DMEM/high glucose medium (31966047, Gibco) supplemented with 10% (v/v) heat-inactivated FBS (10500064, Gibco) and 8 µM 2-Mercaptoethanol (21985023, Gibco), under 5% CO_2_ and 37°C. HEK293T cell line was maintained in high glucose DMEM (31966047, Gibco) supplemented with 10% FBS and 1% Antibiotic-Antimycotic (15240062, Gibco). Cells were maintained in culture by routinely splitting when 70–90% confluence.

### Time-course analysis of gene expression

Cells were seeded at a density of 60.000 cells per cm^2^ in 35 mm diameter cell culture dishes (0003700112, Eppendorf), and allowed to attach for 5 hours. Overnight synchronization was achieved by switching to DMEM with 0.5% FBS. The next day, the medium was replaced with complete medium, and cells were exposed to either high (21%) or low (1%) oxygen concertation conditions, with the culture medium in 1% O_2_ pre-equilibrated overnight in a H35 Hypoxystation (Don Whitley Scientific). Cells were harvested at 6, 12, 24, 36, and 48 hours, lysed in TRIzol Reagent (15596018, ThermoFisher Scientific) and kept at -80^0^C until used for RNA extraction.

### RT-qPCR

RNA extraction and qPCR analysis were performed using standard protocols according to manufacturer’s instructions. Briefly, total RNA was isolated using TRIzol and reverse-transcribed with the High-Capacity RNA-to-cDNA™ Kit (4387406, Applied Biosystems). Quantitative PCR was performed using gene-specific primers and SYBR™ Select Master Mix (4472918, Applied Biosystems) on ViiA™ 7 Real-Time PCR System (ThermoFisher Scientific). Relative RNA expression was assessed by the ΔΔCT method, normalizing to 18S rRNA and Rpl32 mRNA levels. All primer sequences will be shared upon specific request.

### Cell cycle analysis

First the cells were trypsinized and then resuspended in PBS for cell cycle analysis. Then, approximately 10^5^ cells (in 300 ul PBS) were fixed with ice-cold pure ethanol added drop-wise to cell suspension, followed by 3-hour incubation at -20°C. After fixation, cells were stained with 20 µg/ml propidium iodide and 100 µg/ml RNAse A for 30 minutes at 37°C. Samples were subsequently analyzed on a CytoFlex flow cytometer (Beckman Coulter), capturing data from 20,000 events per sample.

### Competitive cell growth assay

Wildtype, miR210-KO, and MIR210HG-KO HEK cells were fluorescently labelled with either dTomato or EGFP through a 48-hour lentiviral infection using the plasmids pUltra-Chili (Plasmid #48687) and pUltra (Plasmid #24129) from Addgene (https://www.addgene.org). To ensure uniform fluorescent labelling, single-cell-derived clones were generated form the GFP/dTomato-expressing cells, which were then further propagated and used for proliferation assays. To assess competitive growth, equal ratios of EGFP-labeled wildtype and dTomato-labeled knockout cells, or the opposite, were co-cultured. After 72 hours, the growth rate of each genotype was determined by flow cytometric analysis using the CytoFLEX system (Beckman Coulter), measuring the percentage of each cell type in the mixed culture.

### RNA-sequencing of HEK cells

RNA-seq was performed by Novogene (UK) Company Limited. The quality of total RNA was assessed using an Agilent 2100 Bioanalyzer. Subsequent library preparation was performed, followed by high-throughput sequencing on the Illumina HiSeq 2500 platform using PE150 mode, generating approximately 20 million reads per sample. Paired-end clean reads were aligned to the reference genome using STAR software (version 2.6.1d). Quantification of reads mapped to each gene was achieved using FeatureCounts (version v1.5.0-p3). Differential expression analysis was carried out using the DESeq2 package (version 1.20.0), with significant genes showing an adjusted P-value < 0.05 designated as differentially expressed.

### RNA sequencing of mouse fibroblasts

Two independent cultures were performed in either 21 or 1% hypoxia during 24h. RNA was extracted with the miRNeasy mini kit (Qiagen) and libraries were constructed for both small RNA-seq and mRNA-seq as previously described (36). Briefly, mRNA-seq was performed from 2 µg of RNA that was first subjected to mRNA selection with Dynabeads® mRNA Purification Kit (Invitrogen). mRNA was fragmented 10 min at 95 °C in RNAseIII buffer (Invitrogen) then adapter-ligated, reverse transcribed and amplified with the reagents from the NEBNext mRNA Library Prep Set for SOLiD. Small RNA-seq was performed from 500 ng RNA with the NEBNext Small RNA Library Prep Set for SOLiD according to manufacturer’s instructions. Both types of amplified libraries were purified on Purelink PCR micro kit (Invitrogen), then subjected to additional PCR rounds with primers from the 5500 W Conversion Primers Kit (Life Technologies). After Agencourt® AMPure® XP beads purification (Beckman Coulter), libraries were size-selected from 150 nt to 250 nt (for RNA-seq) and 105 nt to 130 nt (for small RNA-seq) with the LabChip XT DNA 300 Assay Kit (Caliper Lifesciences), and finally quantified with the Bioanalyzer High Sensitivity DNA Kit (Agilent). Libraries were sequenced on SOLiD 5500XL (Life Technologies) with single-end 50b reads. SOLiD data were analyzed with lifescope v2.5.1, using the small RNA pipeline for miRNA libraries and whole transcriptome pipeline for RNA-seq libraries with default parameters. Annotation files used for production of raw count tables correspond to Refseq Gene model for mRNAs and miRBase for small RNAs. Differential expression analysis was carried out using the DESeq2 package, with significant genes showing an adjusted P-value < 0.05 designated as differentially expressed.

### Transmission Electron Microscopy

Cells were fixed in 2,5% buffered glutaraldehyde followed by post-fixation in 1% osmium tetroxide with 1.5% potassium ferrocyanide in 0.1 M cacodylate buffer. with 1.5% potassium ferrocyanide in 0.1 M cacodylate buffer. Sequential dehydration in 70%, 95% and 100% ethanol were followed by sample embedding in Agar 100 epoxy resin at 60°C for 48 hrs. The ultrathin sections were cut at 80 nm using a diamond knife and double-stained with uranyl acetate and lead citrate. Ultrathin sections were examined using a Morgagni 268 TEM (FEI Company, The Netherlands) at 80 kV. Digital electron micrographs were acquired using a MegaView III CCD using iTEM-SIS software (Olympus, Soft Imaging System GmbH, Germany). The quantification of mitochondria and endoplasmic reticulum relative volumes (V/v) were estimated using a grid superimposed onto electron microscopy images. The images were acquired from 2 sections/sample and 10 different images/section. The relative volumes were estimated by a point-counting method using stereological grids with regularly distributed test-points (at 0,2 µm) generated in iTEM-SIS software (Olympus, Soft Imaging System GmbH, Germany).

### Cellular Bioenergetics Stress

Cells were subjected to Seahorse XF Cell Energy Phenotype Test (103325-100, Agilent) and Seahorse XF Cell Mito Stress Test (103010-100, Agilent), using the Seahorse XFp Analyzer (Agilent, CA, USA) per manufacturer’s instructions, Briefly, cells were seeded at 2×10^5^ cells per well in miniplates for 24h and, the growth medium was switched to the specially formulated assay medium before testing. The Oxygen Consumption Rate (OCR) was measured under both basal and stressed conditions induced by FCCP and oligomycin. Mitochondrial functionality was further interrogated by challenging the cells with sequential addition of oligomycin, FCCP, rotenone, and antimycin A. Data normalization was based on DNA content (Hoechst 33258 staining), and data analysis was performed using Wave 2.6.3 software’s Report Generators.

### Reactive Oxygen Species (ROS) production

ROS production in mouse fibroblasts was quantified using MitoTracker™ Red CM-H2XRos (M7513, ThermoFisher Scientific) and flow cytometry, according to manufacturer guidelines. Briefly, cells were cultured at a density of 2×10^4^ cell/cm^2^ for 24 hours and then exposed for another 24 hours under standard (21% O_2_) or low oxygen (1% O_2_) conditions, as described above. After 48 hours, without changing the oxygen exposure, cells were detached, resuspended in 300 µl DMEM w/o phenol red, and stained with MitoTraker dye at 400 nM for 30 minutes at 37_°_C. Stained cells were then washed and immediately analyzed on a flow cytometer after adding propidium iodide, with data collected from 50,000 events per sample to ensure robust measurements.

### Bioinformatic analysis

Pathway enrichment analysis of DEGs was performed using the Enrichr software tool (37, 38). Briefly, using the Appyter for bulk RNA-seq, DEGs were analyzed utilizing the MSigDB_Hallmark_2020 gene set to identify and cluster significant pathways. Additionally, miRTarBase_2017 gene set was utilized to estimate the odds ratios for microRNAs potentially regulating the input DEG set. The UCSC Genome Browser was used to display the genomic positioning and structure of the MIR210HG locus and the neighboring genes. Long-range chromatin interactions were examined using publicly available Hi-C data available through the 3DIV database (http://3div.kr) (39).

### Statistical analysis

Statistical analysis was performed GraphPad Prism (version 7.05). Unless stated otherwise in the figure legend, the statistical significance of differences was determined by two-way ANOVA, and Tukey test was applied for multiple comparisons. The difference was considered statistically significant when p-value was <0.05. Data are presented as mean ± SD, unless stated otherwise.

## Acknowledgements

This work was supported in part by the NIH R01 CA155332-01A1 (MI), Romanian Academy, Romanian Ministry of Research, Innovation, and Digitization (PN-III-P1-1.1-TE-2019-1893, contract no TE186/2021), and a grant from Romania’s National Recovery and Resilience Plan (PNRR-III-C9-2022-I8-CF186, contract no 760062/23.05.2023). M.B.P. was supported by the Foundation for Cellular and Molecular Medicine. MG was supported by MCID grant PN 23.16.01.01. SBC was supported by Stockholm County Research Council, von Kantzows Foundation, and Kung Gustaf V’s och Drottning Victorias Frimurarestifelse.

## Author contributions

**Conceptualization:** Mircea Ivan; Mihai Bogdan Preda, Alexandrina Burlacu;

**Investigation:** Mihai Bogdan Preda, Evelyn Gabriela Rusu-Nastase, Carmen Alexandra Neculachi, Xiaoling Zhong, Christine Voellenkle, Nathalie M. Mazure, Ovidiu Balacescu, Xiao-Wei Zheng, Mihaela Gherghiceanu, Alexandrina Burlacu;

**Formal Analysis:** Cristina Ivan, Kevin Lebrigand;

**Methodology:** Mihai Bogdan Preda, Fabio Martelli, Bernard Mari, Sergiu-Bogdan Catrina, Alexandrina Burlacu, Mircea Ivan;

**Writing – Original Draft Preparation:** Mircea Ivan, Mihai Bogdan Preda;

**Writing – Review & Editing:** Mircea Ivan; Mihai Bogdan Preda, Ovidiu Balacescu, Mihaela Gherghiceanu, Maya Simionescu, Fabio Martelli, Bernard Mari, Sergiu-Bogdan Catrina, Alexandrina Burlacu;

**Funding Acquisition:** Mihai Bogdan Preda, Fabio Martelli, Sergiu-Bogdan Catrina, Mircea Ivan;

**Supervision:** Mircea Ivan, Mihai Bogdan Preda.

## Data availability

The RNA-seq datasets produced in this study are in the process of being indexed in the specific databases.

## Supporting information

**Supplementary figure 1.**
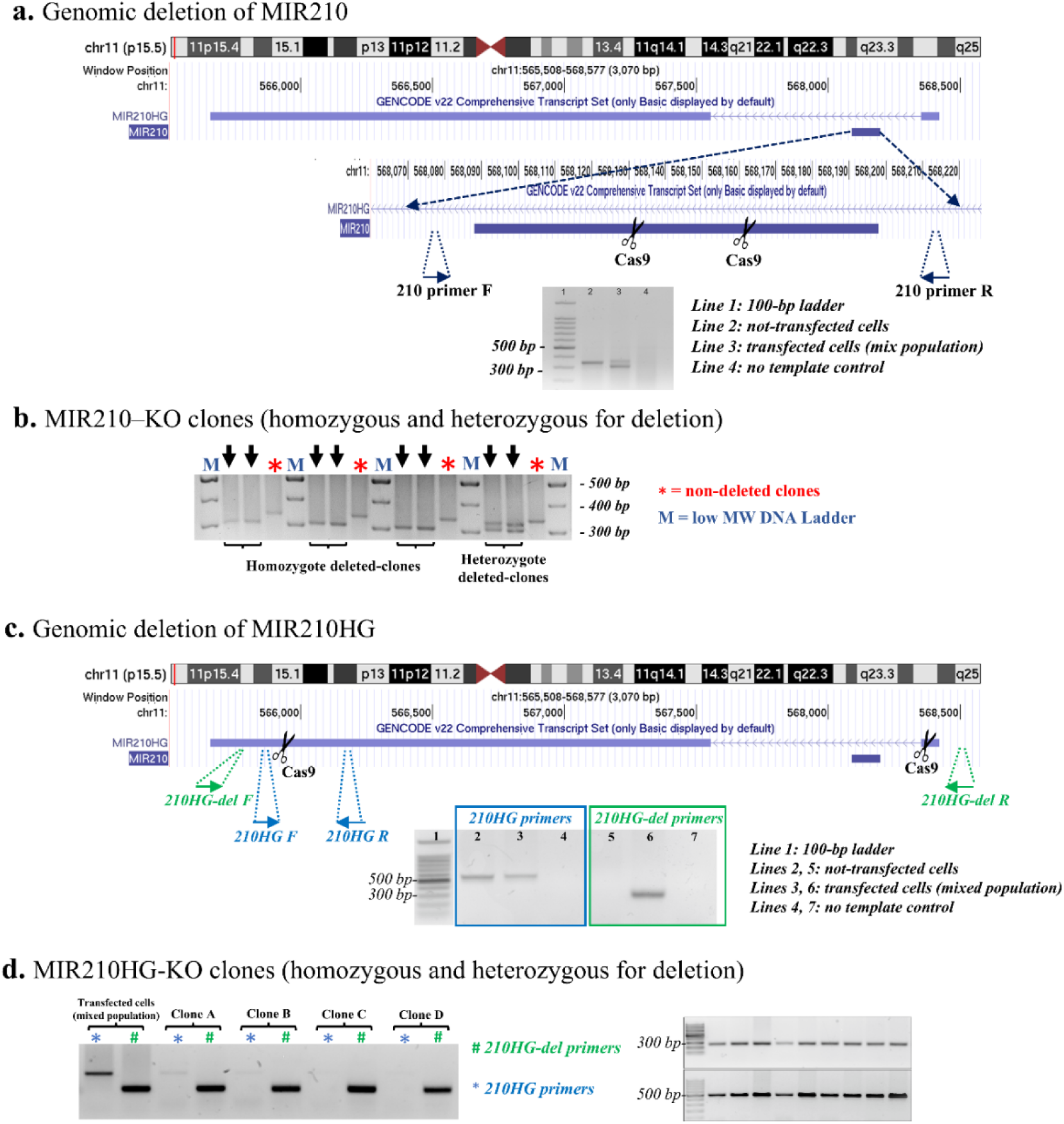
Generation of MIR210-KO and MIR210HG-KO human cells. a) Schematic representation of miR-210 deletion with CRISPR/Cas9 genome editing tool using UCSC Genome Browser to illustrate the genomic location of the miR-210 gene, the ∼ 30 bp fragment deleted and the location of the primers used for validation b) Validation by genomic PCR of different miR210-KO clones that are heterozygous and homozygous for deletion. c) Illustration of MIR210HG deletion on UCSC Genome Browser, showing the ∼2500 nucleotide fragment cut, and the position of the two set of primers used to validate the genomic deletion. d) MIR210HG-KO clones homozygous and heterozygous for genomic deletion.

**Supplementary figure 2.**
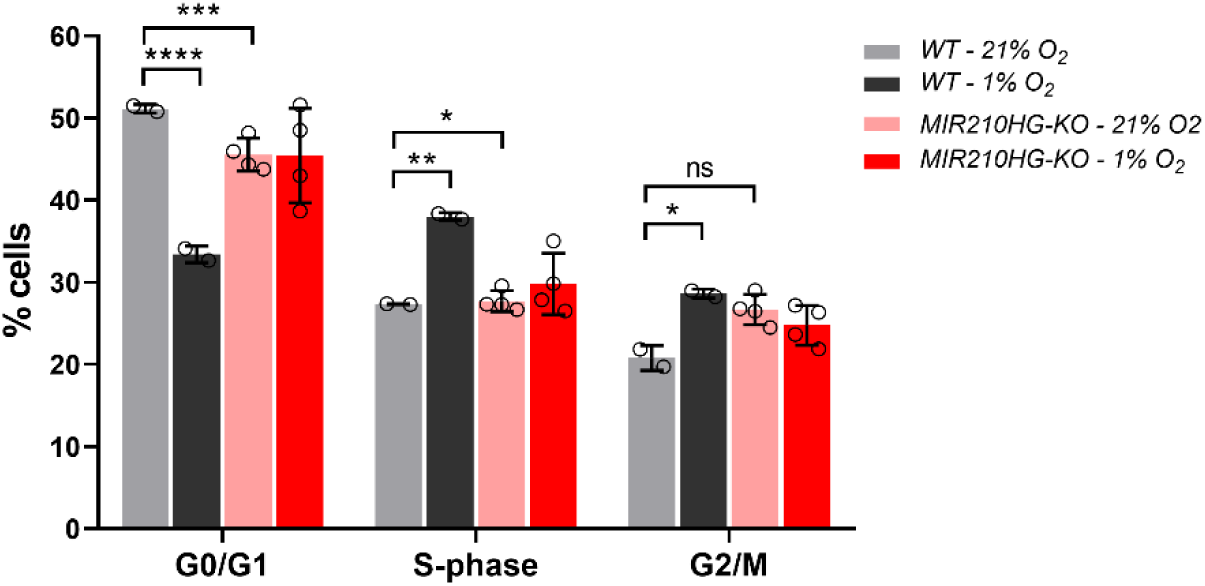
Cell cycle distribution in HEK cells (WT and MIR210HG-KO) under different oxygen levels (two-way ANOVA, *p < 0.05, **p < 0.01, ***p < 0.001, ****p < 0.0001).

**Supplementary figure 3.**
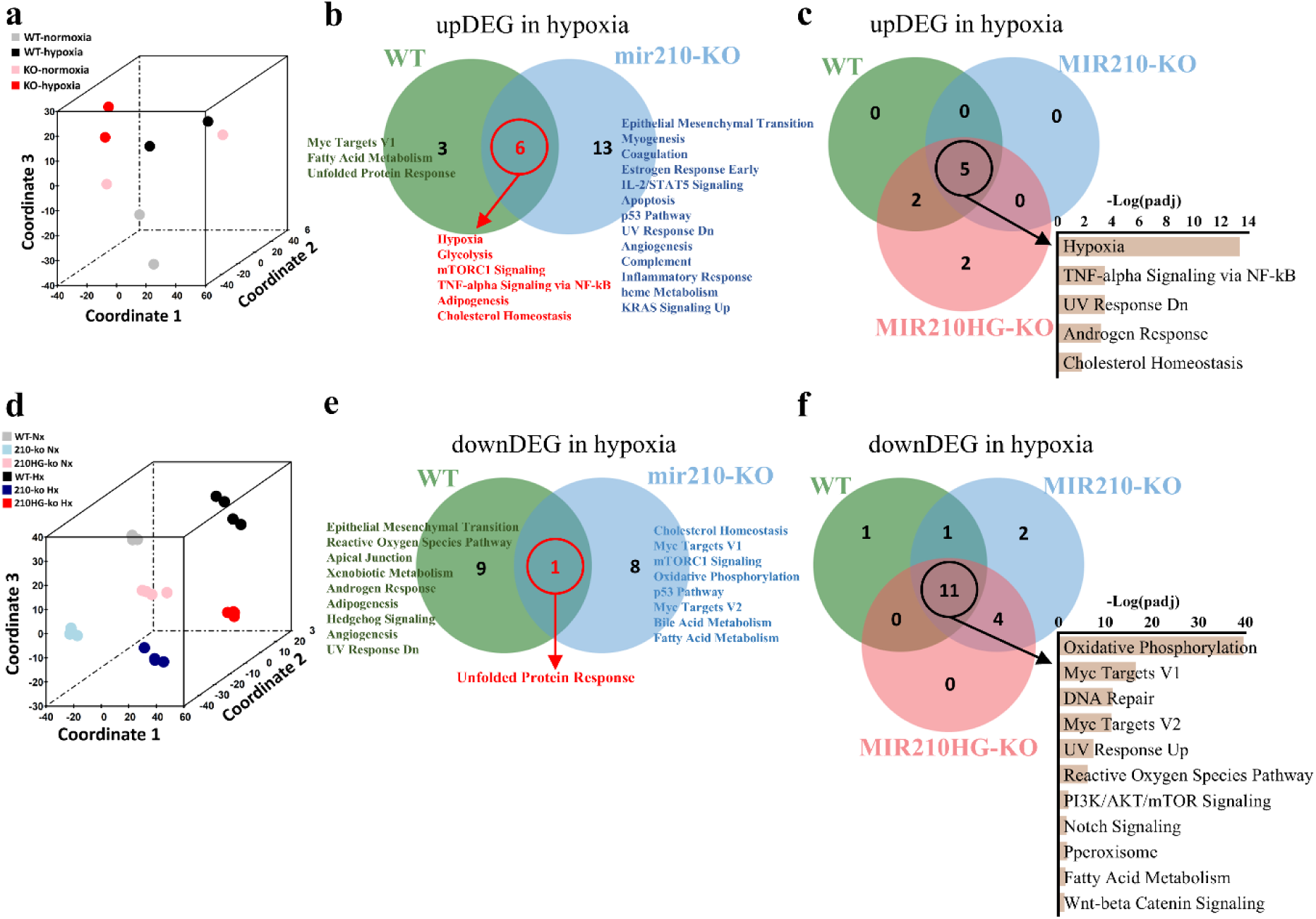
Global transcriptomic changes in miR210-KO and MIR210HG-KO cells. Three-dimension PCA illustrating the overall gene expression structure in wild-type, miR-210-KO or MIR210HG-KO cells under low and high oxygen conditions, in mouse fibroblasts (a) and HEK cells (b). (c-d) Venn diagrams showing distinct and overlapping signaling pathways identified using the Enrichr tool (MSigDB Hallmark 2020 gene set) from upregulated and downregulated differentially expressed genes in mouse fibroblasts exposed to hypoxic versus normoxic conditions. (e-f) Venn diagrams showing distinct and overlapping signaling pathways identified identified from upregulated and downregulated DEGs in WT and KO HEK clones exposed to hypoxic versus normoxic conditions.

**Supplementary figure 4.**
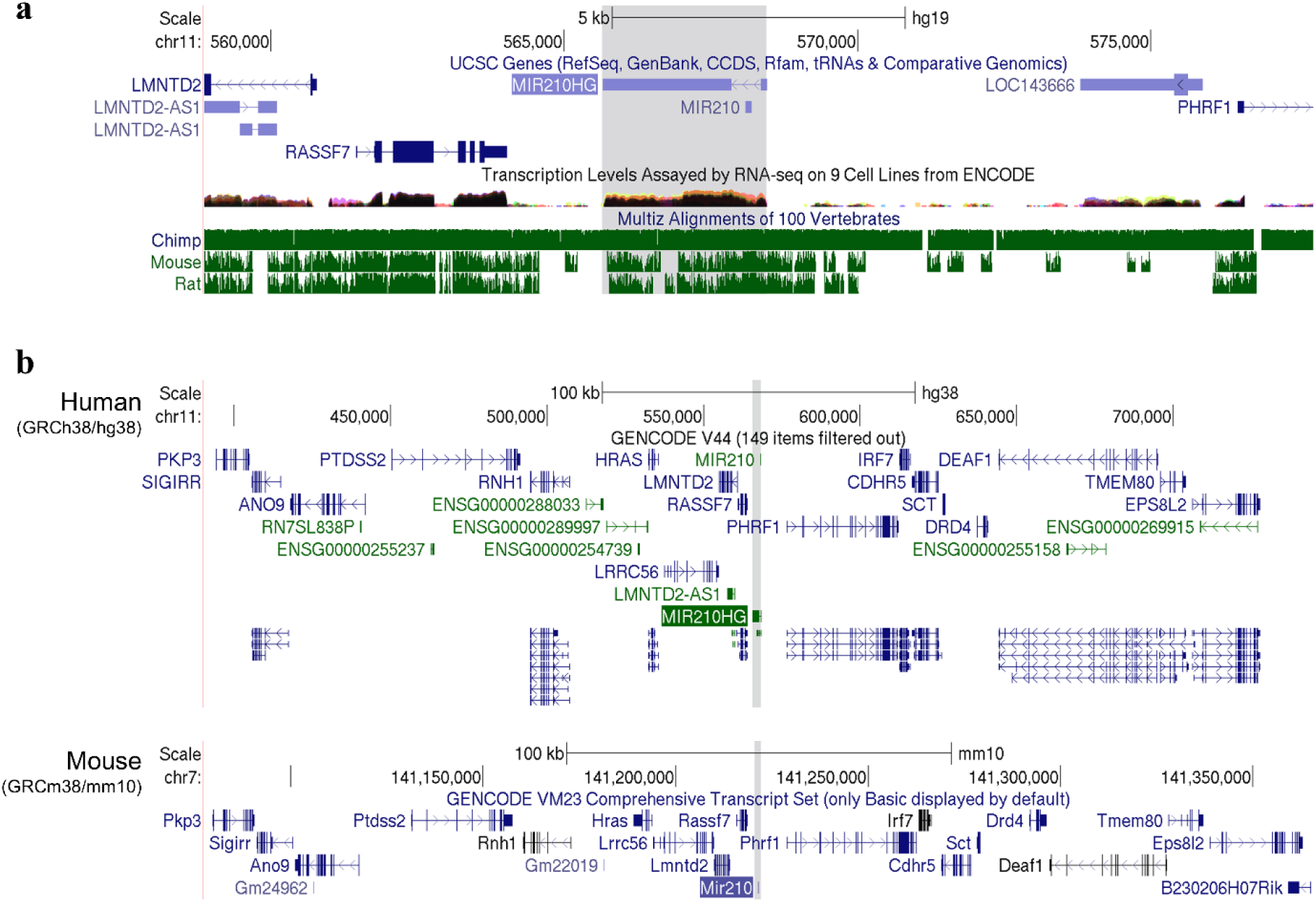
Genomic features of MIR210HG genomic locus. a) Evolutionary conservation between rodents and primates of MIR210HG locus as illustrated using Vertebrate Multiz Alignment & Conservation tracks in UCSC Genome Browser. b) The track of ∼ 300 kb spanning the MIR210 genomic locus (highlighted in gray) which illustrate the protein-coding genes and non-coding RNA genes annotated in GENECODE V46 and visualized using UCSC Genome Browser for human (GRCh38/hg38) and mouse (GRCm38/mm10) genomes.

**Supplementary figure 5.**
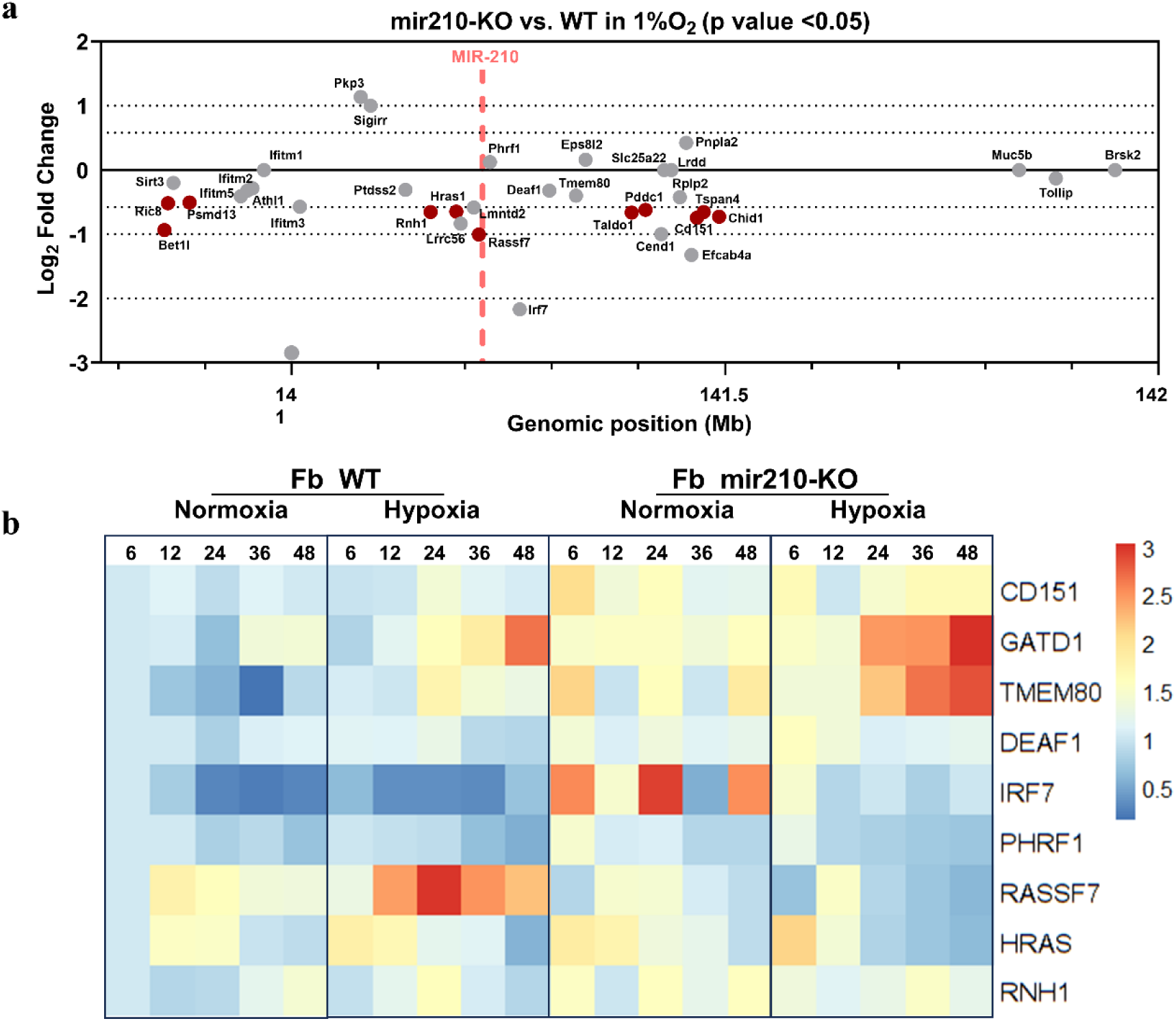
Effect of MIR210 locus deletion on transcriptional gene regulation in mouse fibroblasts. a) Summary of local transcriptional changes upon miR210 deletion in mouse fibroblasts exposed to hypoxia (1% O_2_) for 24 hours. Genes are plotted based on their chromosomal position on the x axis and their differential expression (log2FoldChange) on the y-axis. Genes significantly differentially expressed (p value < 0.05) were name-labelled and marked with red dots. MIR210 genomic locus is highlighted by a vertical dashed line. (b) Heatmap visualization of qRT-PCR analysis of specified neighboring genes at various time points in both 21% and 1% O_2_. Data are presented as fold changes relative to Fb WT Normoxia at 6 hours and are normalized on a per-row color scale for clarity

**Supplementary figure 6.**
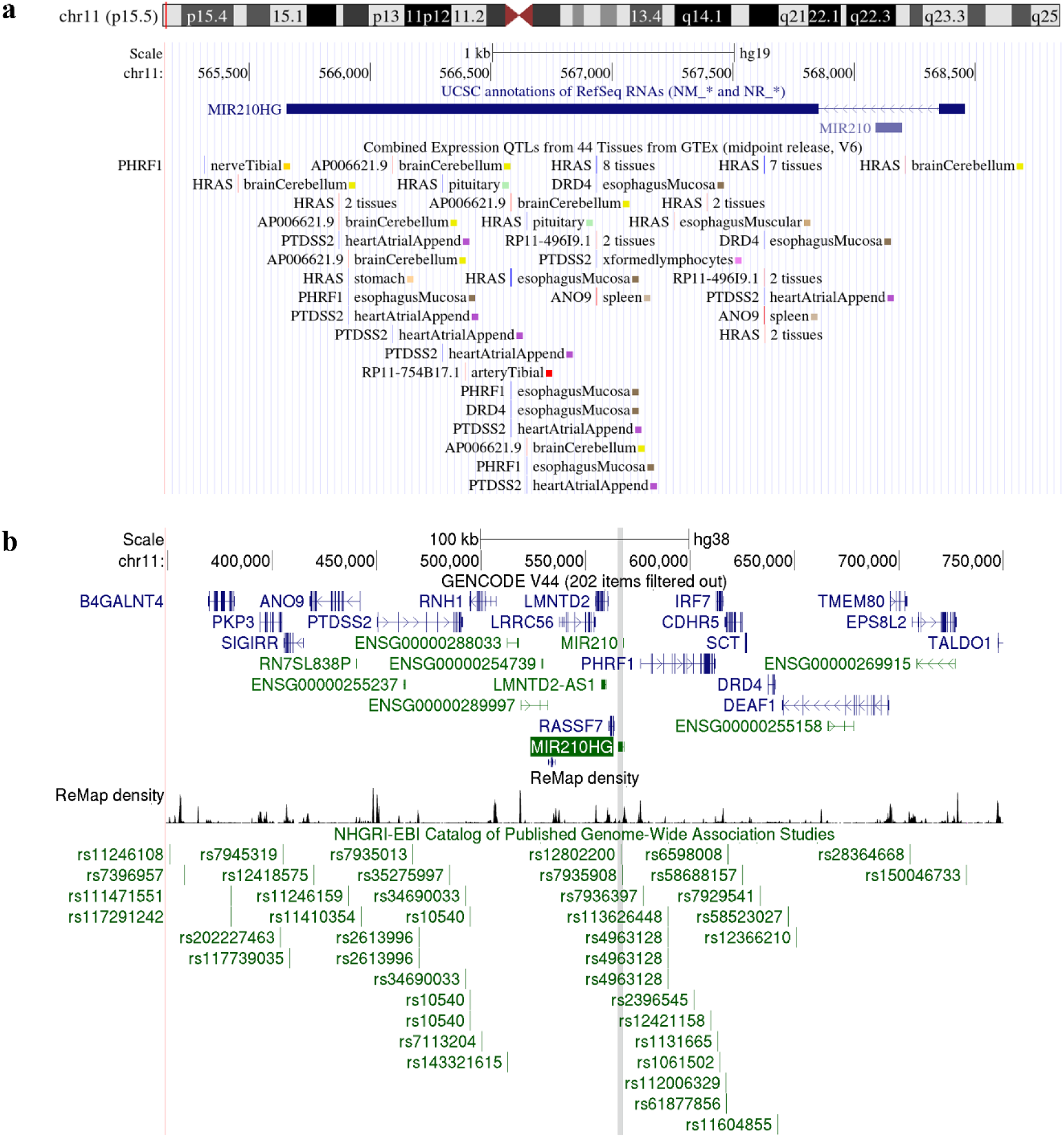
Genomic features of human MIR210HG genomic locus. a) UCSC genome browser track showing genetic variants of MIR210HG locus displayed as gene expression quantitative trait loci within 1MB of gene transcription start sites (cis-eQTLs), that likely affect proximal gene expression in 44 human tissues from the GTEx V6 data release. b) UCSC genome browser track displaying single nucleotide polymorphisms (SNPs) identified by published Genome-Wide Association Studies (GWAS) and collected in the NHGRI-EBI GWAS Catalog.

**Supplementary figure 7.**
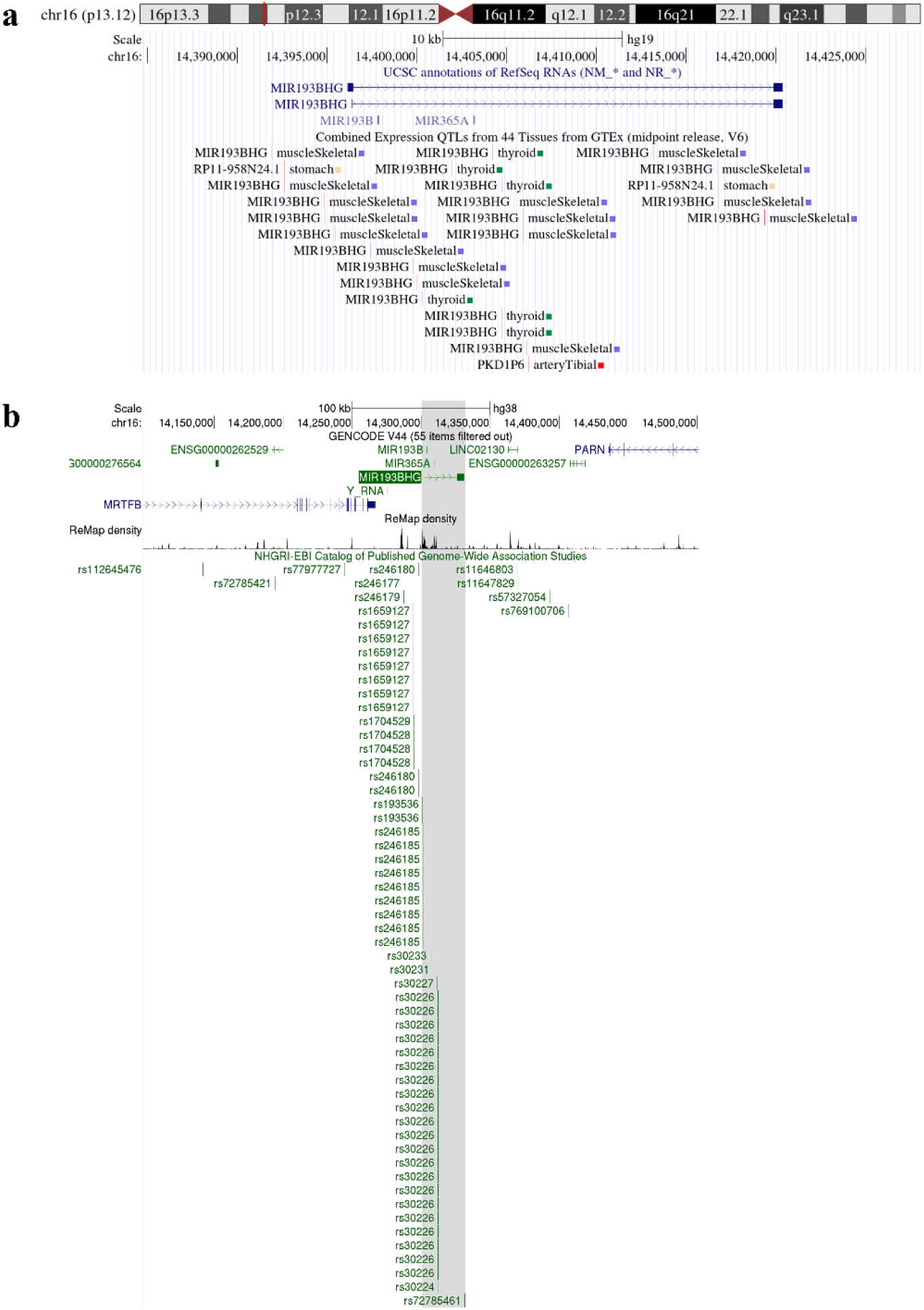
Genomic features of human lincNORS (MIR193BHG) genomic locus. a) UCSC genome browser track showing genetic variants of lincRNORS locus displayed as gene expression quantitative trait loci within 1MB of gene transcription start sites (cis-eQTLs), that likely affect proximal gene expression in 44 human tissues from the GTEx V6 data release. b) UCSC genome browser track displaying single nucleotide polymorphisms (SNPs) identified by published GWAS and collected in the NHGRI-EBI GWAS Catalog.

**Supplementary figure 8.**
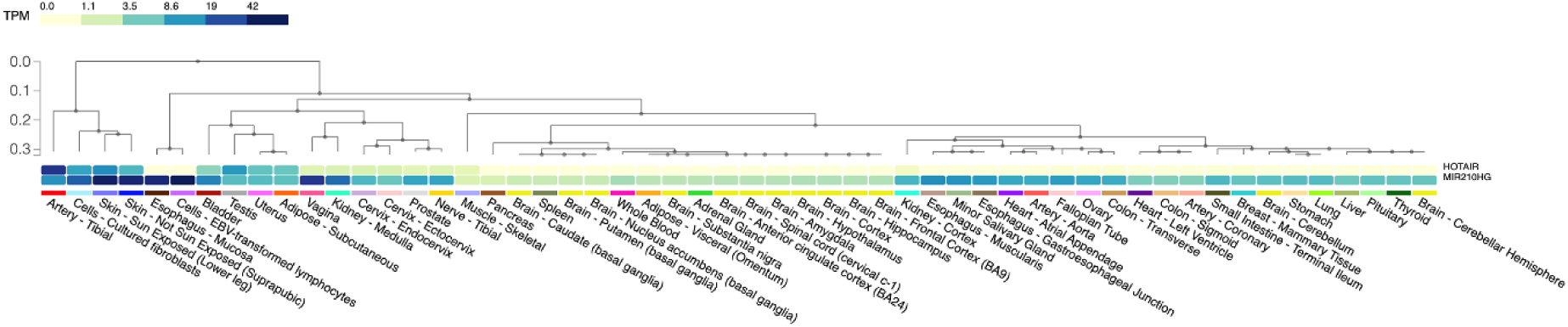
Expression levels (TPM > 0.1) of HOTAIR and MIR210HG lncRNAs across the broad tissue categories in the GTEx database.

